# Cell culture dimensionality influences mesenchymal stem cell fate through cadherin-2 and cadherin-11

**DOI:** 10.1101/744698

**Authors:** Fiona R Passanha, Thomas Geuens, Simon Konig, Clemens A van Blitterswijk, Vanessa LS LaPointe

## Abstract

The acquisition of a specific cell fate is one of the core aims of tissue engineering and regenerative medicine. Significant evidence shows that aggregate cultures have a positive influence on fate decisions, presumably through cell-cell interactions, but little is known about the specific mechanisms. To investigate the difference between cells cultured as a monolayer and as aggregates, we started by looking at cadherin expression, an important protein involved in cell adhesion, during the differentiation of bone marrow-derived human mesenchymal stem cells (hMSCs) in aggregate and monolayer cultures. We observed that proliferating hMSCs in monolayer culture express cadherin-2 and undergo a switch to cadherin-11 over time, which was not evident in the aggregate cultures. By knocking down cadherin-2 and cadherin-11, we found that both cadherins were required for adipogenic differentiation in a monolayer as well as aggregate culture. However, during osteogenic differentiation, low levels of cadherin-2 were found to be favorable for cells cultured as a monolayer and as aggregates, whereas cadherin-11 was dispensable for cells cultured as aggregates. Together, these results provide compelling evidence for the important role that cadherins play in regulating the differentiation of hMSCs and how this is affected by the dimensionality of cell culture.

## INTRODUCTION

For regenerative medicine, cells are typically studied as a monolayer on a flat surface, which does not reflect the conditions most cells experience *in vivo*. Studying cells on flat surfaces strips them of many cell-cell interactions and introduces them to a foreign adherent environment. For example, monolayer culture systems, while perfectly suitable for studying some processes in the dermal epithelium, can fall short for studying developmental processes that do not involve cells in a monolayer (1). As regenerative medicine relies on accurately recapitulating these developmental processes for organ regeneration, culturing or even transplanting aggregates of cells is considered a promising strategy due to enhanced cell-cell interaction and because aggregates more closely mimic the natural environment of a tissue (2, 3). Indeed, cell behavior in aggregate culture systems is different from that in monolayer systems (4), and studies have revealed molecular differences between monolayer and aggregate culture systems (5–7). For example, it has been shown that AKT and mTOR signaling is drastically reduced in aggregate culture systems (5).

Differences in the dimensionality and geometry of cells between monolayer and aggregate culture systems can lead to different cellular responses (8). Aggregate culture systems not only influence the spatial organization of the cell surface receptors engaged in interactions with surrounding cells, but they also induce physical constraints to cells (9). These systems can affect the signal transduction and ultimately influence gene expression and cellular behavior. For example, it has been shown that cadherin-based cell-cell interaction increases compared to integrin-based cell-matrix adhesion in aggregates (10, 11). Adhesion formation on flat surfaces with a focus on integrins is well documented (12, 13), and also studied in aggregate systems, where it was suggested that the type of integrin employed by the cell is differentially specified by the dimensionality of the microenvironment (14). Compared to integrin-based adhesion, less is known about cadherin-based adhesion in aggregate cultures. As regenerative medicine is moving towards a more physiologically relevant setting to improve outcomes, understanding the role of cadherins in the context of cell aggregates is crucial to understanding processes such as stem cell differentiation.

To this end, we investigated the influence of cadherin-mediated, cell-cell interaction in on the differentiation of bone marrow-derived human mesenchymal stem cells (hMSCs) in monolayers and aggregates. hMSCs are nonhematopoietic multipotent cells that have the potential to differentiate into a variety of cell types including, but not limited to, osteoblasts and adipocytes (15). The human cadherin superfamily comprises of over 100 different proteins (16). In an adult, cadherins bind cells with each other in the presence of calcium ions that give form to different tissues. Cadherins not only maintain tissue integrity but also have a role in diverse biological processes such as differentiation, proliferation, polarity, and stem cell maintenance (17–19). The diversity of the cadherin family makes them capable to take on such varied functions (20, 21). Different types of cadherins are expressed in different types of cells; cadherin-2 and cadherin-11 are expressed in mesenchymal-type cells (22, 23). Because they are implicated in cell differentiation (24, 25), we study cadherin-2 and cadherin-11 expression in hMSCs in both aggregate and monolayer cultures, in order to determine whether and how they influence osteogenic and adipogenic differentiation.

We show that cadherin-2 and cadherin-11 maintain their expression over time in aggregate cultures whereas their expression switches from cadherin-2 to cadherin-11 in monolayer culture. Knockdown of cadherin-11 increases cadherin-2 but not vice versa suggesting a one-directional relationship between the cadherins. Functionally, knockdown of cadherin-2 and cadherin-11 led to alterations in the potential of hMSCs to differentiate towards the osteogenic and adipogenic lineages, underscoring the critical role of cell-cell interaction in directing cell fate. While cadherin-2 enhanced mineralization in both culture formats, cadherin-11 interfered with mineralization only in monolayer culture which corroborates that monolayer and aggregate cultures yield different cell behavior.

## RESULTS

### 1. Phenotype identification and trilineage differentiation potential of hMSCs

We first determined the phenotype of hMSCs by assessing them at passage 5 for representative markers by flow cytometry, and by determining their trilineage differentiation potential by using established protocols, all according to the International Society for Cellular Therapy standard (26). The cultured hMSCs were positive for mesenchymal markers CD73 (97.9%), CD90 (99.6%), and CD105 (95.2%) (Fig. S1 A-C) and negative (≤0.014% positive) for hematopoietic markers CD45, CD34, CD11b, CD19, and HLA-DR (Fig. S1D). The cells successfully differentiated into osteogenic, adipogenic, and chondrogenic lineages, which were determined using histological staining with Oil Red O (Fig. S2A), Alizarin Red S, (Fig. S2B) and Safranin O (Fig. S2C), respectively.

### 2. Cadherin-2 expression decreased over time in monolayer cultures

To investigate whether cadherin expression changes over time when hMSCs are cultured as a monolayer, we measured cadherin-2 and cadherin-11 levels at days 1, 5, 7, and 10 (Fig. 1A). In monolayer cultures, hMSCs expressed significantly decreased cadherin-2 levels over time, whereas cadherin-11 levels remained similar across the four time points (Fig. 1 B and C).

**Fig. 1.**
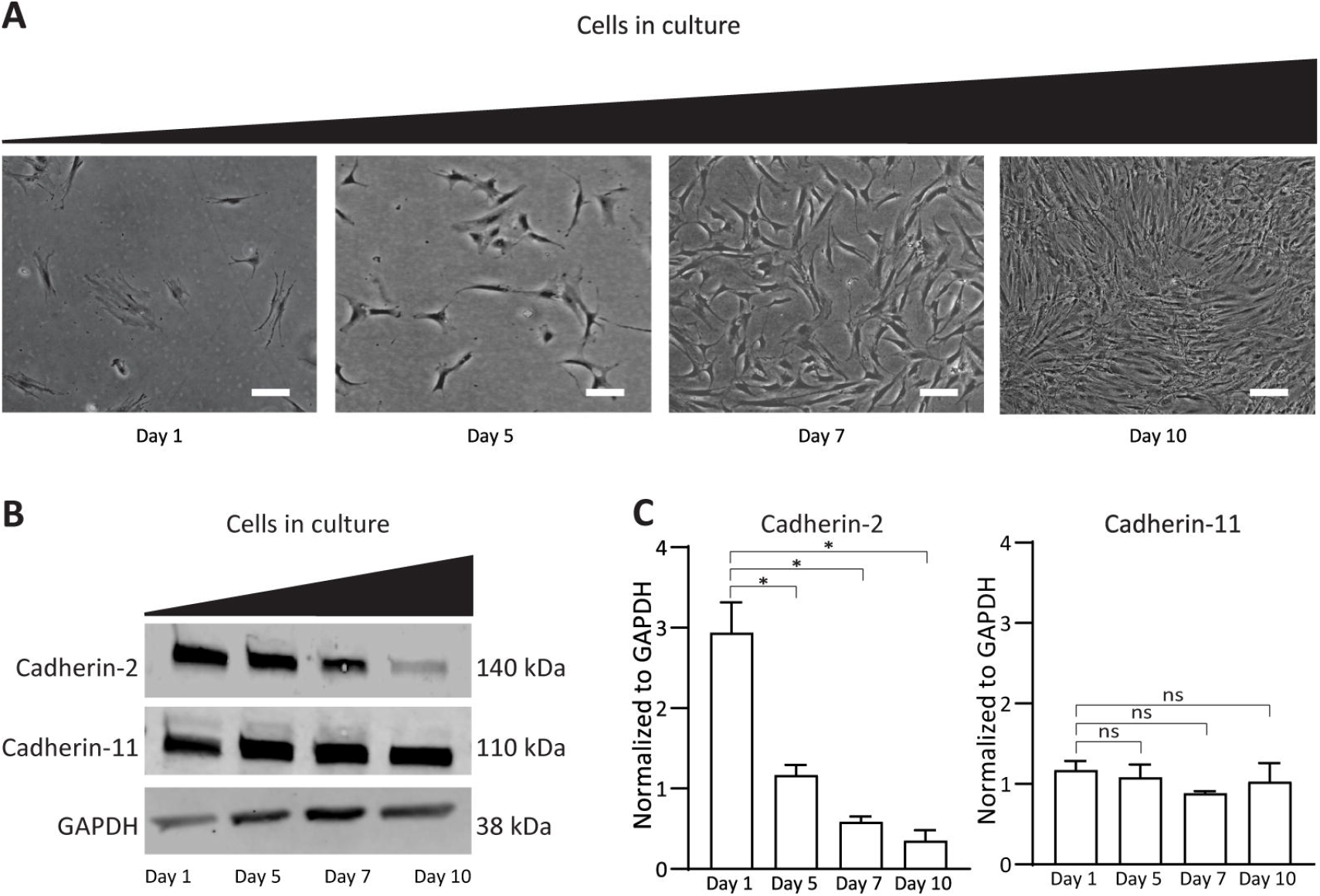
Cadherin-2 levels are inversely related to hMSC cell density in monolayer cultures. hMSCs were seeded at 1 × 10^4^ cells/cm^2^ and evaluated after days 1, 5, 7 and 10. (A) Phase contrast micrographs showed increasing cell density from day 1 to day 10. Scale bars represent 100 μm. (B) Western blots showed decreasing cadherin-2 and stable cadherin-11 expression over time. GADPH is shown as a loading control. (C) Quantification of Western blots normalized to GAPDH showed that cadherin-2 significantly decreased over time while levels of cadherin-11 remained the same. Error bars show ± SD. Data are representative of at least three independent experiments with similar results. Statistics were determined using one-way ANOVA with Holm-Sidak’s test for multiple comparisons: \**p* < 0.01; ns, not significant.

### 3. Cadherin-2 differentially expressed in hMSCs in cultures of different dimensionality

hMSCs were seeded as a monolayer and as aggregates, and cadherin levels were measured after hMSCs attained confluency in monolayer (approximately 5 d) or formed aggregates (24 h) in aggregate cultures. This timepoint, denoted day 0, indicated the start of hMSC differentiation (Fig. 2 A and B). We found that hMSCs in aggregate cultures expressed significantly lower levels of cadherin-2 compared to those cultured in monolayers (Fig. 2 C and D). In comparison, cadherin-11 levels were similar for both monolayer and aggregate cultures at day 0 (Fig. 2 C and D). In monolayer culture on day 1, cadherin-2 was localized to the adherens junctions, while cadherin-11 was not (Fig. 2E). Over time, on day 21, cadherin-2 expression substantially decreased at the adherens junctions, while the expression of cadherin-11 increased (Fig. 2E). The varied distribution of cadherin-11 did not affect the total levels of cadherins in monolayer culture. In aggregate culture, hMSCs expressed cadherin-2 as well as cadherin-11, which was consistent over time (Fig. 2F).

**Fig. 2.**
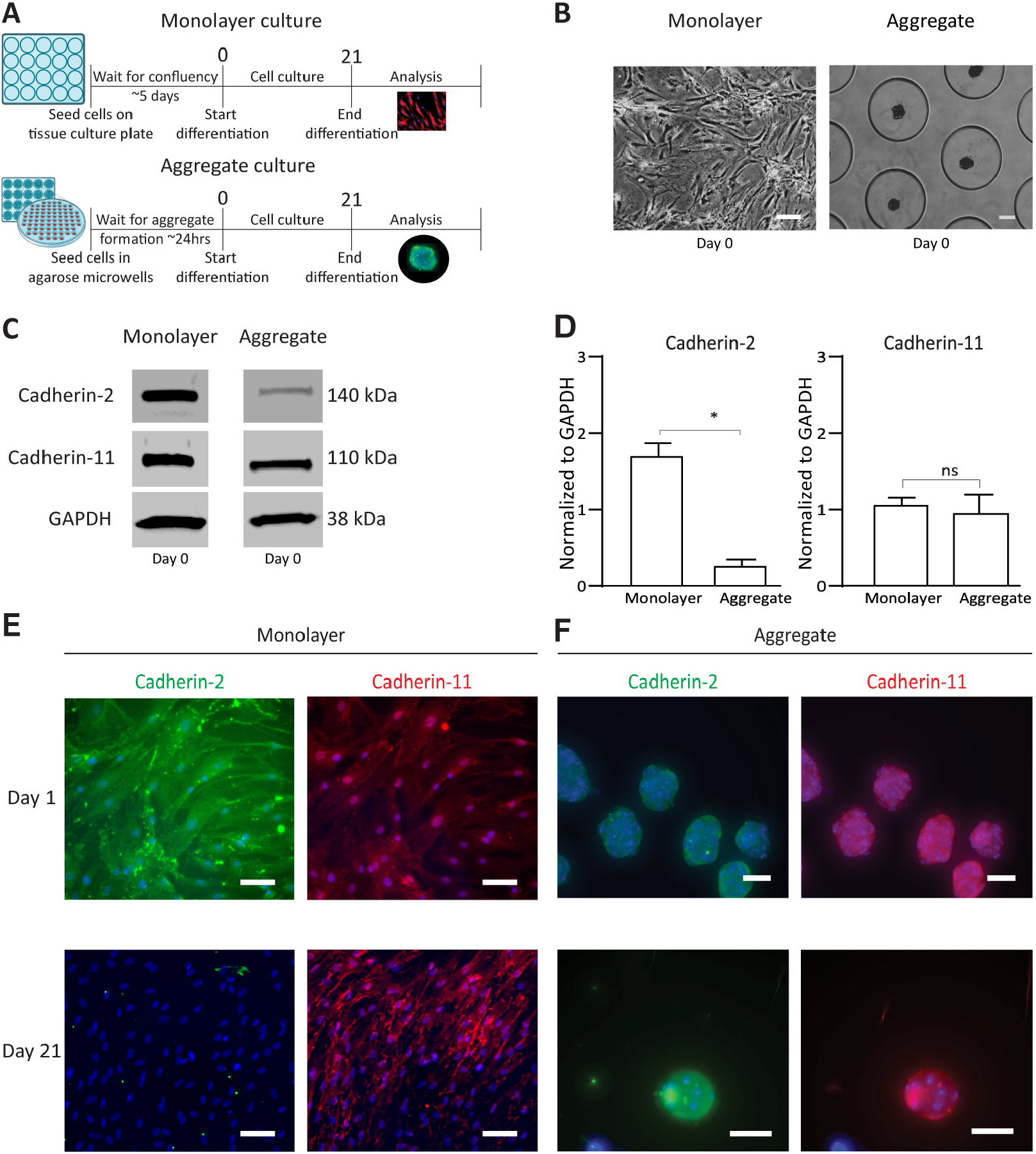
hMSC cultures and expression of cadherin-2 and cadherin-11. (A) Schematic timeline of the cells cultured as a monolayer and as aggregates. (B) Phase contrast micrograph of hMSCs plated as a monolayer and seeded as aggregates in microwells at day 0. Scale bars represent 100 μm. (C) Western blot indicates lower cadherin-2 expression in aggregate culture compared to monolayer culture at day 0, while cadherin-11 expression was similar. GAPDH is shown as a loading control. (D) Quantification of Western blots normalized to GAPDH expression showed a significant decrease in cadherin-2 in aggregate culture compared to monolayer culture while levels of cadherin-11 showed no significant difference. Error bars show ± SD. Data are representative of at least three independent experiments with similar results. Statistics were determined using one-way ANOVA with Holm-Sidak’s test for multiple comparisons: \**p* < 0.01; ns, not significant. (E) Fluorescence micrographs of immunostained hMSCs, cultured as a monolayer with cadherin-2 (green, left column) and cadherin-11 (red, right column) and counterstained with DAPI (blue) showed that cadherin-2 expression decreased between day 1 (top row) and day 21 (bottom row). (F) Fluorescence micrographs of immunostained hMSCs, cultured as aggregates with cadherin-2 (green, left column) and cadherin-11 (red, right column) and counterstained with DAPI (blue) showed that cadherin-2 and cadherin-11 expression were expressed similarly across both timepoints, different from their expression in the monolayer culture.

### 4. Differentiation pathways influenced cadherin levels in monolayer culture but not in aggregate cultures

To investigate whether cadherin expression changed as hMSCs were cultured in differentiation medium, the cells were cultured in osteogenic inductive medium and adipogenic inductive medium and immunostained for cadherin-2 and cadherin-11 at days 1 and 21 for cells cultured as a monolayer, and days 1 and 10 for cells cultured as aggregates. Cadherin expression in cells cultured as a monolayer was influenced by inducing differentiation. On day 1, cadherin-2 was expressed at the adherens junctions (Fig. 3 A and E), which significantly decreased by day 21 in both inductive medium conditions (Fig. 3 B and F). Cadherin-11 at day 1 was expressed in the cytoplasm, which by day 21 was observed at substantially higher levels at the adherens junctions (Fig. 3 D and H). We observed different levels of cadherin-11 expression over time depending on the medium in which the hMSCs were cultured in monolayer (Fig. 3 D and H): osteogenic induction medium resulted in higher levels of cadherin-11 (Fig. 3D) compared to adipogenic inductive medium (Fig. 3H) after 21 days in culture. These data were consistent with Figure 1 and 2, where cadherin-2 decreases and cadherin-11 increases over time in monolayers.

**Fig. 3.**
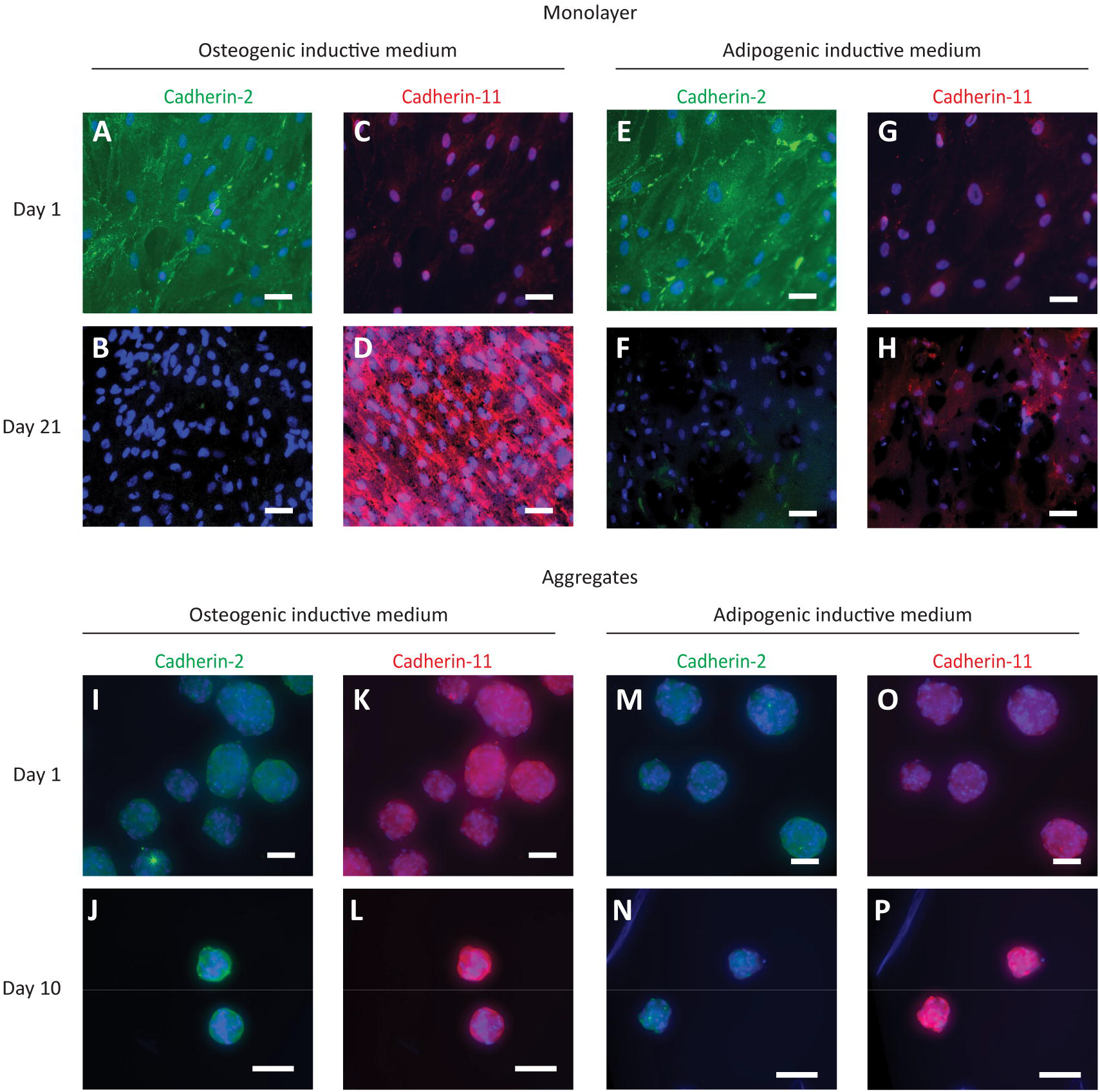
Differentiation medium conditions induced variation in cadherin-2 and cadherin-11 levels in hMSCs. Fluorescence micrographs of hMSCs immunostained with cadherin-2 (green, A, B, E, F, I, J, M, N) and cadherin-11 (red, C, D, G, H, K, L, O, P) and counterstained with DAPI (blue). Cells were grown in a monolayer (A–H) and as aggregates (I–P) in osteogenic (two left columns) or adipogenic (two right columns) induction medium. In monolayers, cadherin-2 expression decreased over 21 days in both differentiation media whereas cadherin-11 expression increased over 21 days in osteogenic induction medium only. In hMSC aggregates, the cadherin expression at day 10 did not differ from the expression at day 1 in either differentiation media. Data are representative of at least three independent experiments with similar results. Scale bars represent 100 μm.

Unlike the situation in monolayer culture, it appeared that cadherin expression in aggregates was uninfluenced by differentiation. In aggregate cultures, cadherin-2 showed similar immunostaining at both time points and with both differentiation media (Fig. 3 I, J, M, N), which was also observed with cadherin-11 (Fig. 3 K, L, O, P); these observations indicate differentiation media did not affect cadherin expression in aggregate cultures.

### 5. Cadherin-11 knockdown increased cadherin-2 expression

To explore whether the changes in cadherin-2 and cadherin-11 expression over 21 days in monolayer culture were interrelated, we performed lentiviral transduction with cadherin-11 shRNA, cadherin-2 shRNA or scrambled shRNA on hMSCs cultured in growth medium as a monolayer. Western blot analysis and qPCR confirmed a knockdown efficiency of 80% for cadherin-11 (Fig. 4 A and B) and 84% for cadherin-2 (Fig. 4 C and D) 7 days after starting selection. The knockdown efficiency was maintained after 21 days in culture (Fig. 4F).

**Fig. 4.**
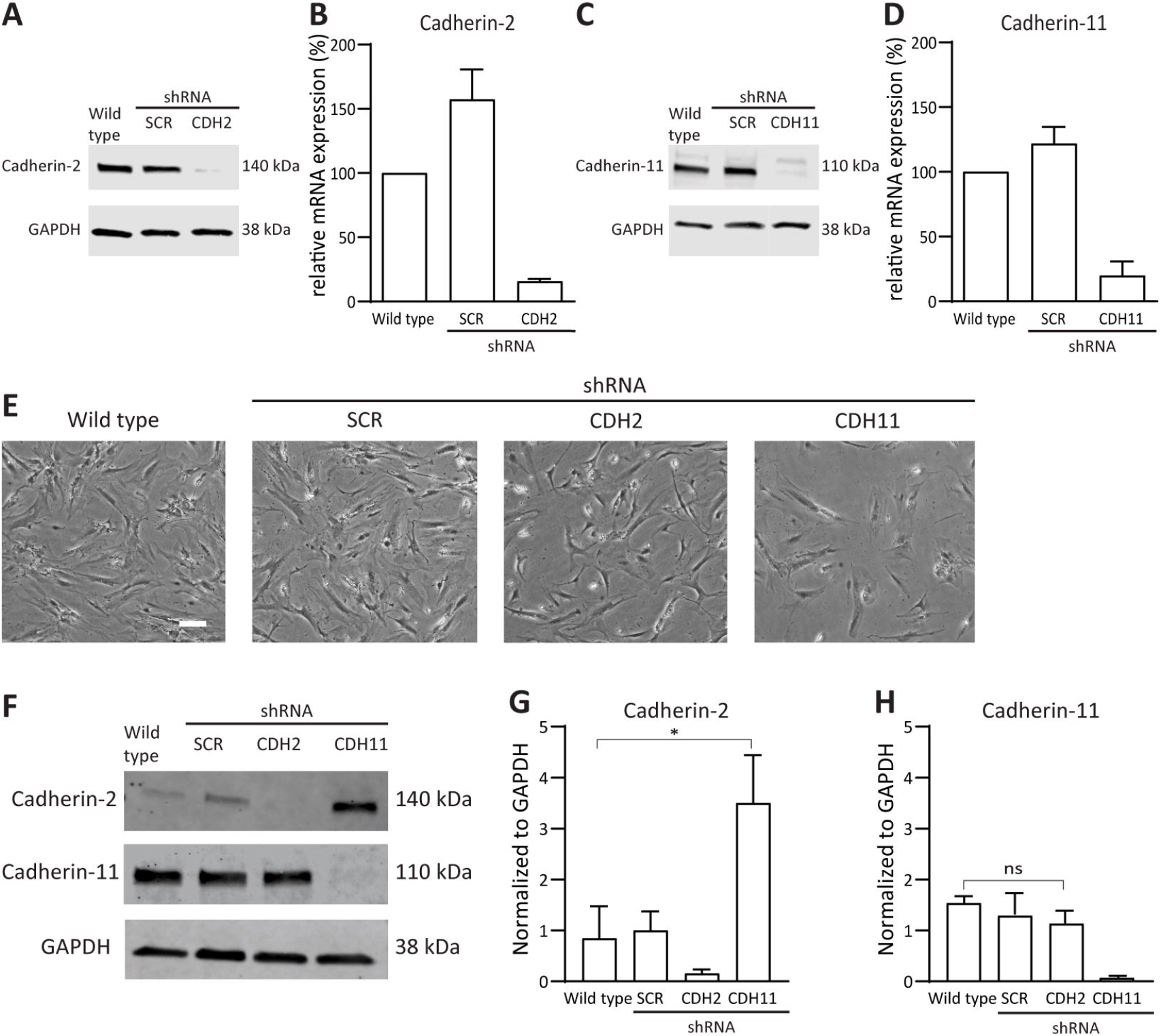
Cadherin-11 knockdown resulted in increased cadherin-2 expression. (A, C) Western blot analysis of cadherin-2 (A) and cadherin-11 (C) showed decreased protein expression by the respective shRNA knockdowns. GAPDH is shown as a loading control. (B, D) Quantification of relative mRNA expression demonstrated 84% knockdown efficiency for cadherin-2 (B) and 80% knockdown efficiency for cadherin-11 (D) compared to controls. (E) Phase contrast micrographs of hMSCs taken 7 days after cadherin-2 and cadherin-11 knockdown revealed morphological changes in cadherin-11–knockdown cells compared to all other conditions. Scale bars represent 100 μm. (F) Western blots showed an upregulation of cadherin-2 in cadherin-11–knockdown cells. In comparison, cadherin-11 expression in cadherin-2 knockdown cells was not affected. GAPDH is shown as a loading control. (G) Quantification of Western blots normalized to GAPDH showed that the increase in cadherin-2 expression in cadherin-11–knockdown cells was significant, while cadherin-11 expression in cadherin-2–knockdown cells was similar to wild-type controls. Error bars show ± SD. Data are representative of at least three independent experiments with similar results. Statistics were determined using one-way ANOVA with Holm-Sidak’s test for multiple comparisons: \**p* < 0.01; ns, not significant. For all panels, SCR indicates hMSCs transduced with a scrambled shRNA knockdown as a negative control.

Morphological changes were observed in cadherin-11 knockdown cells compared to the wild type and the scrambled (Fig. 4E). Surprisingly, we observed that knocking down cadherin-11 caused an upregulation of cadherin-2 expression which was greater than 50% compared to wild-type (Fig. 4 F and G). In comparison, cadherin-11 levels did not change in cadherin-2 knockdown cells (Fig. 4 F and H). These observations indicate that cadherin-2 and cadherin-11 expression in monolayer culture is interrelated and that their expression change over time (Fig. 1–3) seems to be influenced by cadherin-11 but not by cadherin-2.

### 6. Increased mineralized matrix formation in cadherin-2–knockdown hMSCs

Given our observations on differential cadherin expression levels between monolayer and aggregate cultures, we sought to determine whether cadherin-2 and cadherin-11 levels affected osteogenic differentiation. The cadherin-11– and cadherin-2–knockdown cells along with scrambled and wild-type cells were subjected to osteogenic inductive medium for 21 days in both aggregate and monolayer culture. After 21 days in culture, the cells were stained with Alizarin Red S to visualize the mineralized matrix. In both monolayer and aggregate cultures, cadherin-2–knockdown cells showed enhanced mineralized matrix formation compared to the controls (Fig. 5 A and B). Cadherin-11–knockdown cells showed decreased mineralized matrix in monolayer cultures (Fig. 5A) but similar mineralized matrix in aggregate cultures (Fig. 5B) compared to the controls. These observations indicate that low cadherin-2 level enhances the deposition of mineralized matrix during osteogenic differentiation in both culture formats, whereas cadherin-11 expression is important for osteogenic differentiation in monolayers only.

**Fig. 5.**
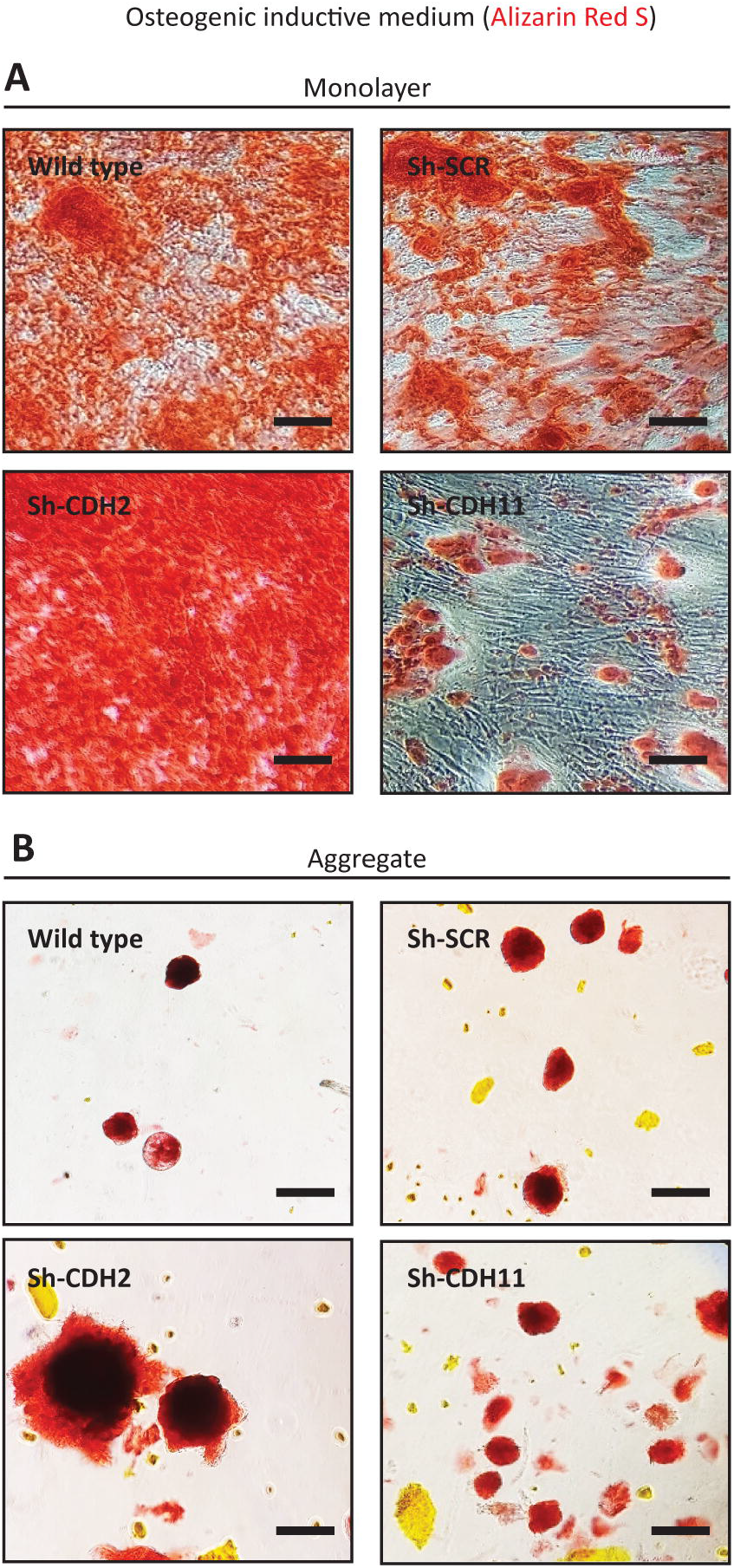
Increased mineralized matrix formation with cadherin-2 knockdown. Brightfield micrographs of hMSCs after 21 days in osteogenic inductive medium as monolayer (A) and aggregate (B) culture. Cadherin-2 and cadherin-11 knockdowns were induced into the osteogenic lineage for 21 days and stained with Alizarin Red S to visualize mineralized matrix. In both monolayer and aggregate culture, cadherin-2 knockdown (sh-CDH2) resulted in increased mineralized matrix formation compared to wild-type controls. In comparison, cadherin-11 knockdown (sh-CDH11) reduced mineralized matrix in monolayer but had no effect on matrix mineralization in aggregate culture compared to wild-type controls. Mineralized matrix following knockdown with a scrambled shRNA (Sh-SCR) is shown as a negative control. Data are representative of at least three independent experiments with similar results. Scale bars represent 100 μm.

### 7. Disrupted adipogenic differentiation in cadherin-2– and cadherin-11–knockdown hMSCs

Next, we investigated how cadherin-2 and cadherin-11 expression influenced adipogenic differentiation. The cadherin-11– and cadherin-2–knockdown cells along with scrambled and wild-type cells were subjected to adipogenic inductive medium for 21 days in both aggregate and monolayer culture. After 21 days in culture, the cells were stained with Oil Red O to visualize the degree of lipid accumulation. Lipid accumulation was substantially reduced in cadherin-2- and cadherin-11-knockdown hMSCs in both monolayer and aggregate culture compared to scrambled and wild-type hMSCs (Fig. 6 A and B). These observations indicate that both cadherin-2 and cadherin-11 are critical for adipogenic differentiation regardless of the dimensionality of the cell culture.

**Fig. 6.**
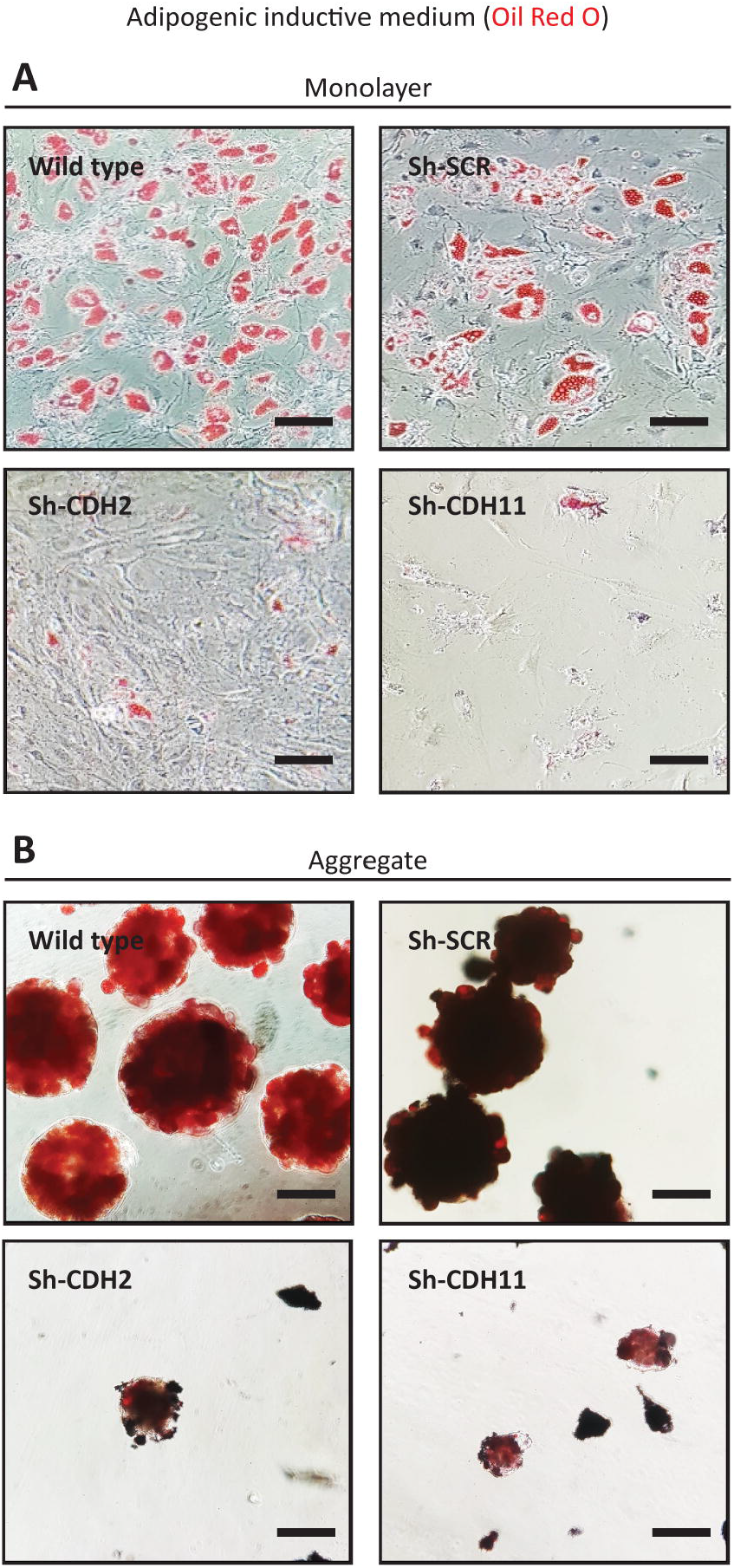
Disrupted adipogenic differentiation potential in hMSCs with cadherin-2 and cadherin-11 knockdowns. Brightfield micrographs of hMSCs after 21 days in adipogenic inductive medium as monolayer (A) and aggregate (B) culture. Cadherin-2 (Sh-CDH2) and cadherin-11 (Sh-CDH11) knockdowns were induced into the adipogenic lineage for 21 days and stained with Oil Red O to visualize lipid accumulation. In both monolayer and aggregate culture, cadherin-2 and cadherin-11 knockdown resulted in a reduced lipid accumulation compared to wild-type controls, indicating diminished differentiation potential. Lipid accumulation following knockdown with a scrambled shRNA (Sh-SCR) is shown as a negative control. Data are representative of at least three independent experiments with similar results. Scale bars represent 100 μm.

## DISCUSSION

The influence of the cell–material interface on cell fate has been an area of significant research in regenerative medicine, but comparatively little is currently known about cell-cell interactions. Furthermore, there is significant evidence that for regenerative medicine, three-dimensional aggregate cultures of hMSCs positively influence fate decisions (27, 28), pointing to a role for cell-cell contact. Our aim was to look at cadherin expression and develop a better understanding of their role in cell fate decisions in monolayer and aggregate cultures.

We began by examining the levels of cadherin-2 and cadherin-11 expressed by hMSCs. During early embryogenesis, mesenchymal tissues have higher expression of cadherin-11 and a comparatively lower cadherin-2 expression (29) which correlates with what we observed during early aggregate formation (Fig. 2). However, in monolayer culture, we found that cadherin-2 expression was greater than cadherin-11 and that these levels decreased with higher cell density over time, while cadherin-11 did not change (Fig. 2). Mesenchymal condensation or mesenchymal cellular aggregation is a critical step for organogenesis. High levels of cadherin-2 have been reported prior to mesenchymal cellular aggregation (30, 31). The ease of condensation in aggregate cultures compared to monolayer culture could explain the differential cadherin expression. However, previous studies on mesenchymal stem cells have shown an increase in both cadherins with increasing cell density in culture (32). Here we have shown that high protein levels of cadherin-2 correspond with a low density of hMSCs (Fig. 1) and do not necessarily indicate engagement of cadherin-2 molecules with their counterparts on the surface of neighboring cells. Our findings suggest that hMSCs have a mechanism that helps regulate cadherin-2 based on the proximity of one cell to another. Specifically, the mechanism downregulates cadherin-2 expression as the distance between cells decreases. The striking observation was that even with ~85% knockdown of cadherin-2 and cadherin-11, hMSCs were able to aggregate, which suggests other, additional mechanisms of cell-cell adhesion. This observation contrasts with a previous study where cadherin-2-depleted cells did not form aggregates (33), which could indicate that even a small amount of cadherin-2 is sufficient for initiation of cell aggregation.

Our observations and other studies have indicated that osteogenesis is favorably affected by low levels of cadherin-2. Knockdown of 84% mRNA enhanced mineralized matrix formation (Fig. 5). In rat MSCs, overexpression of cadherin-2 has been reported to inhibit osteogenesis (34). However, the conditional deletion of cadherin-2 in mice has a negative effect on bone growth because it reduces β-catenin abundance at cell-cell contacts (35). Germline cadherin-2 null mutation is lethal and hence does not allow for the precise understanding of the biological role of cadherin-2 at different stages of osteogenesis *in vivo* (36).

Similar to mRNA knockdown of cadherin-2, an 80% mRNA knockdown of cadherin-11 interfered with the differentiation ability of hMSCs. It reduced mineralized matrix deposition in monolayer culture, in agreement with studies in which cadherin-11–null mutant mice have reduced bone density (37). The effect of the knockdowns on mineralized matrix deposition was more apparent in monolayer culture compared to aggregate cultures, again revealing the important differences cells experience in these two environments. For adipogenesis, both cadherin-2 and cadherin-11 expression were critical (Fig. 6). Cadherin switching during differentiation has been demonstrated in previous studies (18). Interestingly, we observed a switch from cadherin-2 to cadherin-11 expression in monolayer but not in aggregate cultures (Fig. 1). The upregulation of cadherin-11 and the downregulation of cadherin-2 were more pronounced when hMSCs were in osteogenic inductive media. This agrees with the finding that cadherin-11 is highly expressed in cells of the osteogenic lineage (38). However, in aggregate culture, we found no evidence of a switch in cadherin expression during differentiation, as a co-expression of the cadherin-2 and cadherin-11 was observed throughout. These results indicate that cells express different molecules in interactions with their neighbours in monolayer and aggregate cultures.

While cadherin-2 and cadherin-11 are independent members of the cadherin superfamily, our studies and others indicate a relationship between them. For example, the function of cadherin-11 may be in part compensated by cadherin-2 and *vice versa* in mouse models (35, 39). Knockdown of cadherin-11 significantly increased cadherin-2 levels (Fig. 4 F and G). However, this relationship seems to be one-directional, as cadherin-2 knockdown did not affect cadherin-11 levels (Fig. 4 F and H). The upregulation of cadherin-2 as a result of cadherin-11 depletion could be the reason for reduced mineralized matrix deposition in monolayer cultures (Fig. 5A), as overexpression of cadherin-2 inhibits osteogenesis (34). Whether cadherin-11 depletion also upregulates cadherin-2 expression in aggregate cultures remains to be determined. We hypothesize that the upregulation of cadherin-2 in aggregate cultures may not be to the same deleterious levels, because cadherin-2 expression was much lower in aggregates compared to monolayers (Fig. 1) and mineralized matrix deposition was not affected (Fig. 5B). Here we cannot dismiss the possibility of compensation by other cadherins as well. It has previously been shown that cells compensated for the loss of cadherin-2 by upregulation of other cadherins but were unable to compensate for cadherin-2 functionally (40).

Our studies focused on two cadherins, whereas the entire superfamily of 100 cadherins play important roles in tissue integrity and may functionally compensate for one another. Indeed, disrupting cadherins might have a subtle effect on tissue integrity. Conditional knockdown of cadherin-1 or cadherin-3 did not disrupt epidermal integrity in the mouse skin while germline depletion of both cadherin-1 and cadherin-3 caused defects (41). Furthermore, the upregulation of cadherin-2 can functionally compensate for a lack of cadherin-1 in embryonic stem cells (42). These data suggest that individual cadherin species may not be solely responsible for tissue integrity because of compensation by other cadherins. Together, our findings indicate that the nature of cadherin-mediated adhesion is crucial for cell fate determination. Just by looking at two cadherins, we see that cadherins, although very influential in monolayer culture might not be as critical for aggregate culture. Hence, for regenerative medicine, incorporating cadherins in biomaterial design could be beneficial (43).

## MATERIALS AND METHODS

### Monolayer cell culture

Bone marrow-derived hMSCs (PromoCell) were obtained at passage 1. Mycoplasma testing was performed using the mycoplasma detection kit from BD Biosciences. The cells were maintained in growth medium composed of minimal essential medium (MEM α, Gibco) supplemented with 10% fetal bovine serum (FBS). The medium was changed every 2 days, and the cells were maintained at 37°C in 5% CO2 in a humidified incubator. Upon reaching 80% confluence, cells were detached by incubating with 0.05% trypsin-EDTA and replated for continuous passage. The cells were used at passage 5 for all experiments.

### Microwell formation

Agarose microwell arrays were prepared as previously described (44) and inserted into 12-well plates. Each microwell array contained 450 microwells with a diameter of 400 μm.

### Aggregate formation

To form hMSC aggregates, 180,000 cells in a 400 μL suspension in growth medium were seeded into one microwell array. The plate was subsequently centrifuged at 300 ×*g* for 5 min to allow the cells to settle into the microwells, after which an additional 2 mL of growth medium was added to each well. The cells clustered spontaneously within 24 h to form aggregates of approximately 400 cells in each microwell. The medium was changed every 2 days.

### Induction and evaluation of adipogenic and osteogenic differentiation

hMSCs in monolayer culture were expanded to confluency prior to differentiation, while cells in aggregate culture were grown in growth medium for 24 h. All supplements are from Sigma-Aldrich unless mentioned otherwise. To induce osteogenic differentiation, medium was changed every second day with osteogenic inductive medium composed of growth medium supplemented with 0.01 M β-glycerophosphate, 0.2 mM ascorbic acid, and 0.1 μM dexamethasone. To induce adipogenic differentiation, the medium was changed every second day with adipogenic inductive medium composed of Dulbecco’s modified Eagle medium (high glucose, no sodium pyruvate; Gibco) supplemented with 10% FBS, 40 mM indomethacin, 83 mM 3-isobutyl-1-methylxanthine, 10 mg/mL insulin and 0.1 mM dexamethasone. The cultures were maintained for 21 days, after which they were evaluated by Alizarin Red S or Oil Red O staining. Cells were washed twice with phosphate-buffered saline (PBS), fixed in 4% (wt/vol) paraformaldehyde for 15 min at ambient temperature, and washed three times with distilled water. For hMSCs cultured in osteogenic inductive medium, the mineralized extracellular matrix was stained with 2% (wt/vol) Alizarin Red S (VWR) solution in distilled water (pH 4.2) for 15 min. For hMSCs cultured in adipogenic inductive medium, the intracellular lipid accumulation was stained with 0.2% (wt/vol) Oil Red O solution in 60% isopropanol for 15 min.

### shRNA lentiviral transduction

pLKO.1 plasmids containing short hairpin RNA (shRNA) sequences targeting cadherin-11 or cadherin-2 were obtained from Sigma-Aldrich together with a scrambled negative control. These plasmids were co-transfected with third generation lentiviral packaging and envelope vectors; pMD2.G, pRSV-Rev and pMDLg/pRRE (Addgene plasmid #12259, #12253 and #12251, respectively, which were gifts from Didier Trono (45)), into HEK-293T cells using PEIpro (VWR) transfection reagent. The viral supernatant used to transduce hMSCs was collected 24 h after transfection. Forty-eight hours after transduction, positive cells were selected with 2 μg/ml puromycin dihydrochloride (Sigma-Aldrich) in growth media for 5 days. The knockdown efficiency was assessed by qPCR and Western blot after 7 days.

### Immunofluorescence

hMSCs were washed twice with PBS and fixed in 4% paraformaldehyde for 15 min at ambient temperature. For monolayer cultures, fixed cells were washed three times with PBS for 10 min, permeabilized with 0.2% Triton X-100 for 20 min, washed three more times, blocked in 1% goat serum in PBS for 1 h, and incubated with primary antibodies in 0.1% goat serum at 4°C overnight. The cells were washed three times, incubated with secondary antibodies in 0.1% goat serum for 2 h at ambient temperature, and the nuclei were counterstained with DAPI (0.1 μg/mL) for 10 min. For aggregate cultures, fixed cells were washed three times with PBS by centrifugation at 300 × *g* for 5 min, permeabilized with 0.5% Triton X-100 for 1 h, washed three more times, blocked in 1% goat serum in PBS for 1 h, and incubated with primary antibodies in 0.1% goat serum at 4°C overnight with gentle shaking. The cells were washed three times, incubated with secondary antibodies and DAPI in 0.1% goat serum at 4°C overnight with gentle shaking. All samples were mounted in ProLong Gold (Thermo Fisher Scientific), and fluorescence images were acquired on a Nikon E600 inverted microscope. Primary antibodies used were rabbit polyclonal anti-cadherin-11 and mouse monoclonal anti-cadherin-2 (3B9) (both 1:100; Thermo Fisher Scientific). Secondary antibodies used were goat anti-mouse Alexa Fluor 488 and goat anti-rabbit Alexa Fluor 647 (both 1:500; Thermo Fisher Scientific).

### Western blotting

hMSCs were lysed in radioimmunoprecipitation assay (RIPA) buffer supplemented with protease inhibitor (Sigma-Aldrich) and phosphatase inhibitor (Thermo Fisher Scientific). Total protein concentration was measured using the Pierce BCA protein assay kit (Thermo Fisher Scientific). 10 μg of total protein lysate was supplemented with Laemmli buffer, reduced with 5% 2-mercaptoethanol (Sigma-Aldrich) and separated on 4–15% TGX gel (Bio-Rad) followed by transferring to a PVDF membrane (Bio-Rad) using the wet transfer method. Membranes were blocked in 5% milk in Tris-buffered saline (TBS) with 0.1% Tween-20 for 60 min before overnight incubation at 4°C with primary antibodies: rabbit polyclonal anti-cadherin-11, mouse monoclonal anti-cadherin-2 (3B9, both 1:100, Thermo Fisher Scientific) and mouse monoclonal anti-GAPDH (D4C6R, 1:1000, Cell Signaling Technology). Secondary antibodies used were IRDye 680RD goat anti-mouse IgG and IRDye 800CW donkey antirabbit IgG (both 1:15,000; LI-COR Biotechnology). Membranes were imaged on an Odyssey infrared imaging system (LI-COR Biotechnology). Band intensities were determined by quantifying the mean pixel gray values using the ImageJ 1.52b software. Mean pixel gray values were measured in a rectangular region of interest and normalized to GAPDH.

### qPCR

hMSCs were lysed with Trizol (Thermo Fisher Scientific) followed by chloroform phase separation. The aqueous phase was diluted with 70% ethanol in a 1:1 ratio, loaded on an RNA microcolumn (RNeasy mini kit, Qiagen) and RNA extraction was performed according to the manufacturer’s protocol. Subsequently, 600 μg total RNA was converted to cDNA with iScript cDNA Synthesis Kit (Bio-Rad). Real-time PCR was performed in 20 μL reactions using the iQ SYBR Green Supermix (Bio-Rad) and a Real-Time PCR Detection System (Bio-Rad). The cycling conditions were as follows: enzyme activation at 95 °C for 3 min followed by 38 cycles at 95 °C for 12 s and at 58 °C for 30 s. Specific transcripts were detected with the primers listed in Supplementary Table S1 following evaluation for their amplification efficiency.

### Statistical analysis

Statistics were determined using one-way ANOVA with Holm-Sidak’s test for multiple comparisons. *p* values < 0.01 were considered significant.

## ACKNOWLEDGMENTS

We are grateful to Hang Nguyen for critical review of the manuscript. This work was supported by the AO Foundation, Switzerland (S-14-29L) and the Dutch Province of Limburg.

