## Supplementary Figures for "Cell culture dimensionality influences mesenchymal stem cell fate through cadherin-2 and cadherin-11"

**Supplementary Information**

**S1. Characterization of hMSC markers.**

Flow cytometry analysis indicated that the cultured hMSCs expressed surface markers including CD73 (A), CD90 (B), and CD105 (C). Hematopoietic markers CD45, CD34, CD11b, CD19, and HLA-DR (D) considered to be negative mesenchymal markers, were not expressed in hMSCs. The white area indicates the isotype control and black area shows specific signal from the antibody.

***S2. hMSC multipotency***

hMSC multipotency was confirmed by histochemical staining of osteogenic differentiation by Alizarin Red S (A), adipogenic differentiation by Oil Red O (B), and chondrogenic differentiation by Safranin O (C) staining.

### Supplementary figure 1

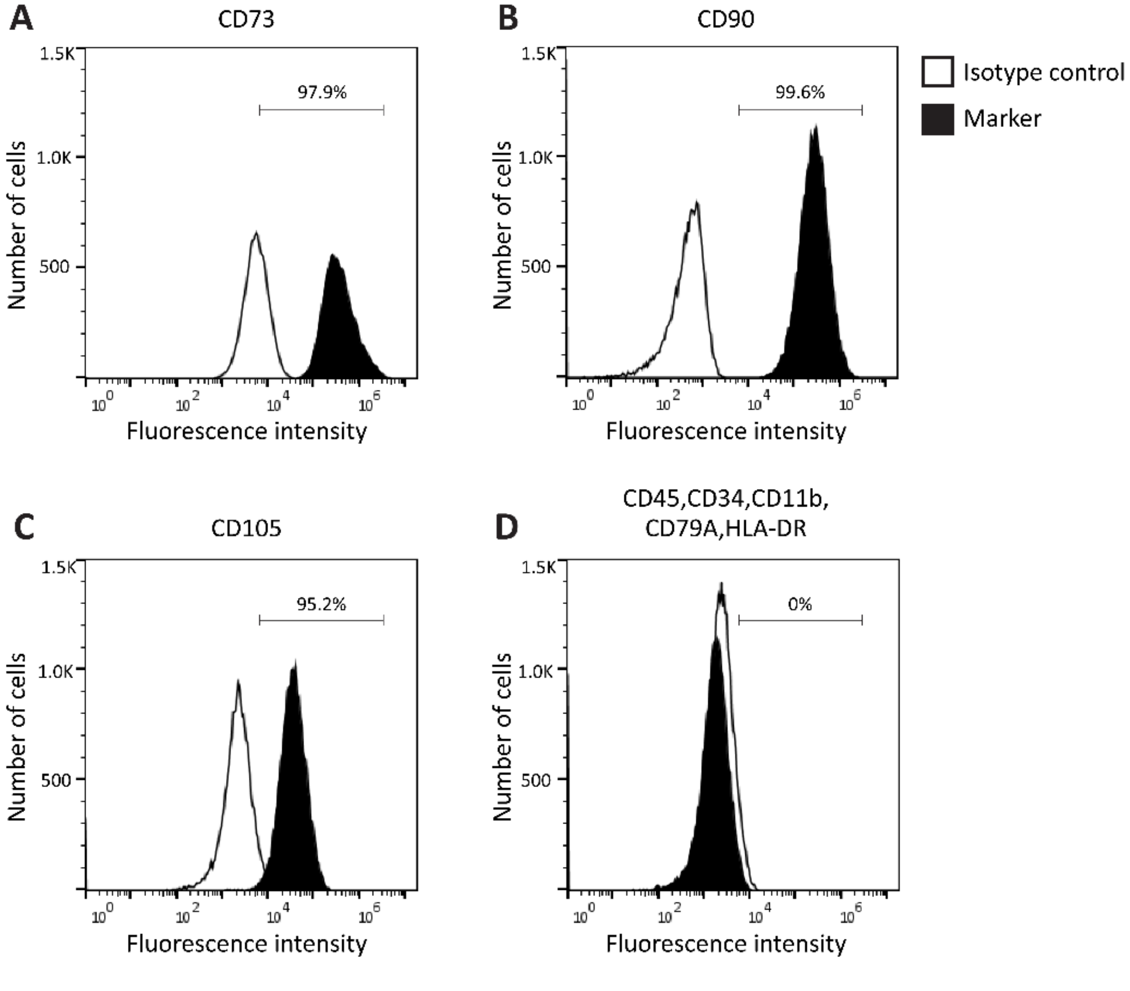

### Supplementary figure 2

**A** Osteogenic inductive medium (Alizarin Red S)

**B** Adipogenic inductive medium (Oil Red O)

**C** Chondrogenic inductive medium (Safranin O)

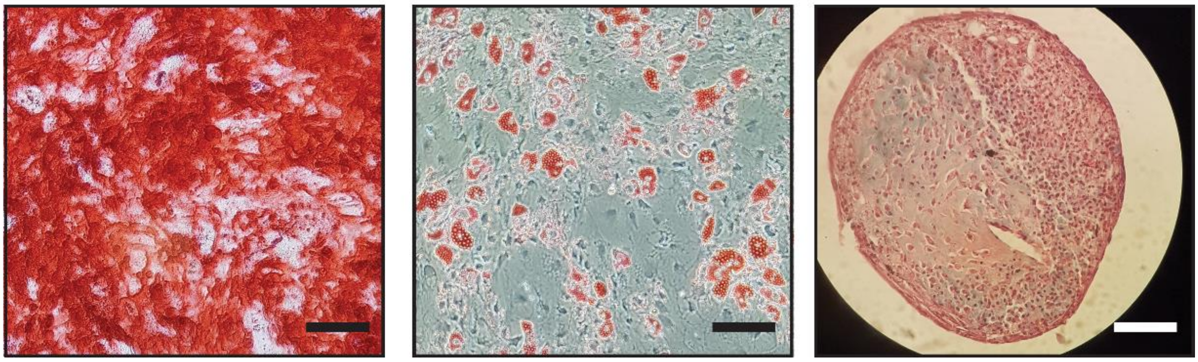
