## Supplementary table for "Cell culture dimensionality influences mesenchymal stem cell fate through cadherin-2 and cadherin-11"

**Table S1**: Primer sequences for RT-PCR

| Gene | Forward primer sequence (5’-3’) | Reverse primer sequence (5’-3’) |
|---|---|---|
| *Cadherin-11 (CDH11)* | AGAGGTCCAATGTGGGAACG | GGTTGTCCTTCGAGGATACTGT |
| *Cadherin-2 (CDH2)* | AGCCAACCTTAACTGAGGAGT | GGCAAGTTGATTGGAGGGATG |
| *Glyceraldehyde-3-phosphate dehydrogenase (GAPDH)* | CTGGGCTACACTGAGCACC | AAGTGGTCGTTGAGGGCAATG |
